# Decoding the Brain: Neural Representation and the Limits of Multivariate Pattern Analysis in Cognitive Neuroscience

**DOI:** 10.1101/127233

**Authors:** J. Brendan Ritchie, David Michael Kaplan, Colin Klein

## Abstract

Since its introduction, multivariate pattern analysis (MVPA), or “neural decoding”, has transformed the field of cognitive neuroscience. Underlying its influence is a crucial inference, which we call the Decoder’s Dictum: if information can be decoded from patterns of neural activity, then this provides strong evidence about what information those patterns represent. Although the Dictum is a widely held and well-motivated principle in decoding research, it has received scant philosophical attention. We critically evaluate the Dictum, arguing that it is false: decodability is a poor guide for revealing the content of neural representations. However, we also suggest how the Dictum can be improved on, in order to better justify inferences about neural representation using MVPA.

## 1. Introduction

Since its introduction, multivariate pattern analysis (MVPA)—or informally, neural ‘decoding’—has had a transformative influence on cognitive neuroscience. Methodologically, it is a veritable multi-tool that provides a unified approach for analyzing data from cellular recordings, fMRI, EEG, and MEG, which can also be paired with computational modeling and behavioral paradigms (Kriegeskorte *et al*. [2008]). Theoretically, it is often presented as a means for investigating the structure and content of the brain’s population code, thereby unifying psychological and neuroscientific explanations while predicting behavioral performance (Haxby *et al*. [2014]; Kriegeskorte and Kievet [2013]). More ambitiously still, decoding methods are advertised as a means of ‘reading’ the brain and ‘listening’ in on the mind (Haynes and Rees [2006]; Norman *et al*. [2006]).

Underlying these bold pronouncements is a crucial inference, which we call the Decoder’s Dictum:

> If information can be decoded from patterns of neural activity, then this provides strong evidence about what information those patterns represent.

The Decoder’s Dictum should interest philosophers for two reasons. First, a central philosophical issue with neuroimaging is its use in ‘reverse inferences’ about mental function (Poldrack [2006]; Klein [2010]). The Decoder’s Dictum is a similar but more nuanced form of inference, so it deserves careful scrutiny. Second, decoding results are some of the most compelling in cognitive neuroscience, and offer a wellspring of findings that philosophers may want to tap into when defending theoretical claims about the architecture of the mind and brain.^1^ It is therefore worth clarifying what decoding can really show.

We argue that the Decoder’s Dictum is false. The Dictum is underwritten by the idea that uncovering information in neural activity patterns, using ‘biologically plausible’ MVPA methods that are similar to the decoding procedures of the brain, is sufficient to show that this information is neurally represented and functionally exploitable. However, as we are typically ignorant of the precise information exploited by these methods, we cannot infer that the information decoded is the same information the brain exploits. Thus decodability is not (by itself) a reliable guide to neural representation. Our goal is not to reprimand neuroscientists for how they currently employ and interpret MVPA. Rather, what follows will clarify the conditions under which decoding could provide evidence about neural representation.

By analogy, consider research on brain-machine interface (BMI) systems, which use decoding to generate control signals for computer cursors or prosthetic limbs (Hatsopolous and Donoghue [2009]). Largely because of BMI’s engineering and translational objectives, however, little attention is paid to the biological plausibility of decoding methods. Consequently, BMI research does not involve inferences about neural function based on decodability. We believe that, epistemically, decoding in cognitive neuroscience is typically no better off than in BMI research, and so forms a thin basis for drawing inferences about neural representation.

Our focus is on how MVPA is used to investigate neural representations. Since talk of representation is itself philosophically contentious, we assume a relatively lightweight notion that is consistent with usage in the relevant sectors of neuroscience: a representation is any internal state of a complex system that serves as a vehicle for informational content and plays a functional role within the system based on the information that it carries (Bechtel [1998]).^2^ As we shall see, some researchers talk of decoding mental representations. We assume they have in mind at least the notion of (distributed) internal representation we have articulated, so our arguments apply to their claims as well.

We focus on neural representations that take the form of population codes. A population code represents information through distributed patterns of activity occurring across a number of neurons. In typical population coding models, each individual neuron exhibits a distribution of responses over some set of inputs, and for any given input, the joint or combined response across the entire neural population encodes information about the input parameters (Pouget *et al*. [2000]).

Our critique of the Dictum will take some setup. In section 2, we provide a brief introduction to decoding methods. In section 3, we argue that the Dictum is false: the presence of decodable information in patterns of neural activity does not show that the brain represents that information. Section 4 expands on this argument by considering possible objections. In section 5, we suggest a way to move beyond the Dictum. Section 6 concludes the paper.

## 2. A Brief Primer On Neural Decoding: Method, Application, And Interpretation

We begin by providing a brief introduction to basic decoding methods and their interpretation. We focus primarily on research that has used MVPA with fMRI to investigate the visual system. There are three reasons for this narrow focus. First, decoding research on vision is largely responsible for popularizing MVPA. Second, it has also driven many of the methodological innovations in the field. Third, it is instructive because we have a detailed understanding of the functional organization of many visual brain regions along with good psychophysics (Haxby [2012]). Thus, if the Dictum is viable at all, it should apply to decoding research on the visual system.

### 2.1 What is MVPA?

Multivariate pattern analysis (MVPA) is a set of general methods for revealing patterns in neural data.^3^ It is useful to separate MVPA into three distinct stages (Mur *et al*. [2009]; Norman *et al*. [2006]), which we will illustrate via a simple (hypothetical) fMRI experiment. In this experiment, fMRI BOLD responses are measured while participants view two gratings of different orientations over a number of trials (Figure 1A). The goal of the experiment is to test whether the activity patterns elicited in response to the two stimulus conditions can be differentiated.

The first step of analysis, pattern measurement, involves collecting neuroimaging data that reflects condition-dependent patterns of activity. This step has a number of components, including performing the actual recordings and preprocessing of the activity-dependent signal. Our example uses fMRI, but other techniques (for example, EEG, MEG, or cellular recordings) could also be employed. As in all fMRI experiments, we must make certain assumptions about the connection between the recorded signals and underlying neural activity.^4^ Nevertheless, the end result is the same: a set of data consisting of multiple distinct measurements of activity occurring during each experimental condition.

**Fig. 1.**
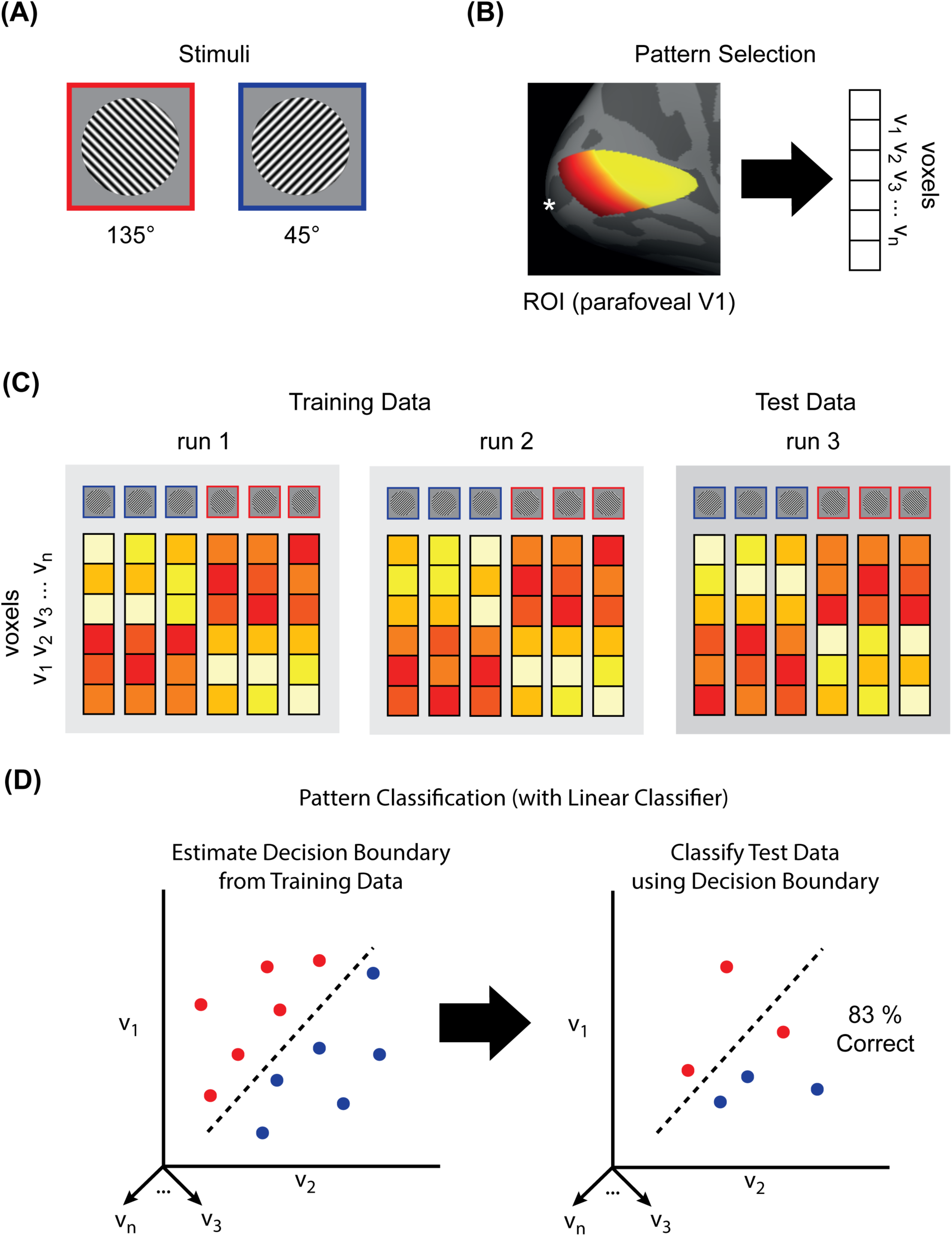

The second step, pattern selection, involves focusing in on a subset of the measured signals for further analysis. With fMRI, this involves a subset of all voxels or a ‘region of interest’ (ROI). ROIs can be defined anatomically (using connectivity patterns or architectonic criteria) and/or defined functionally (using neural response profiles or more traditional univariate fMRI analyses). Pattern selection also depends on experimenter goals and recording technique. In our experiment (Figure 1B) the ROI is parafoveal primary visual cortex (V1), defined anatomically (Benson *et al*. [2012]).

The third, and crucial, step is pattern classification. Pattern classification allows one to measure the discriminability of different patterns in multivariate data. For example, in our experiment we want to see if the patterns of BOLD activity in parafoveal V1 for our two stimulus conditions can be distinguished (Figure 1C). A number of classification methods are available. The simplest is to divide the data in half for each stimulus condition and compute the within- and between-class correlations of the patterns (Haxby *et al*. [2001]). If the patterns are discriminable, the within-class correlation should be higher.

A more powerful (and widely used) technique employs machine learning classifiers, which treat each element of the patterns of interest (e.g., each voxel) as a separate dimension, or ‘feature’, in a high-dimensional space. Assuming our ROI includes *N* voxels, then each trial-wise stimulus presentation elicits a pattern that occupies a point in an *N*-dimensional neural activation space. The goal of the classifiers is to find a way to transform this high-dimensional space into one where the voxel patterns associated with each condition are separable by a decision boundary (Figure 1D).

Although a rich variety of classifiers are available, usually simple linear classifiers are used for MVPA because they provide a principled means of estimating a linear boundary between classes in activation space. To avoid overfitting, the decision boundary is estimated for a subset of the data designated as ‘training’ data, and the classifier is subsequently ‘tested’ on the remaining data (Figure 1D). The classifier assigns condition labels for the training data based on the position of the activity patterns relative to the decision boundary. The performance of the classifier is then a function of the accuracy of its label assignments (for example, % correct; Figure 1D). Training and testing is done multiple times, with each data partition taking its turn as the testing data, and the performance of the classifier is then averaged across iterations. If the mean classifier performance is statistically better than chance, the patterns for the different conditions are considered to be discriminable. Although applications are typically far more complex than what we have presented here, at root all decoding analyses make use of either correlations or machine learning classifiers.

### 2.2 The informational benefits of MVPA

Before we turn to the Dictum, it is worth considering the advantages of MVPA over more traditional univariate analysis methods. To do this we adapt a distinction from Kriegeskorte and Bandettini ([2007]) between activation-based and information-based analyses of neuroimaging data. Activation-based analysis involves spatially averaging activity across all voxels within a given ROI, yielding a single measure of overall regional activation to correlate with the tested conditions. By contrast, information-based analysis looks for a statistical dependency between experimental conditions and the detailed local spatiotemporal activity patterns distributed across the set of individual voxels comprising the ROI (see, for example, Haxby *et al*. [2014]; Tong and Pratte [2012]). Hence, what distinguishes the two approaches is whether or not they are sensitive to spatial patterns in fMRI data. Information-based approaches are so-called because they are sensitive to information contained in these spatial patterns. In contrast, the spatial averaging at the heart of activation-based analyses obscures this information.

All MVPA methods are information-based. Consequently, whatever the status of the Dictum, MVPA decoding holds an advantage over most univariate methods because it offers more spatially sensitive dependent measures. Demonstrating that information is present in activity patterns is also likely to have greater functional significance given the widely held assumption that the brain is an information-processing system that uses population coding to implement its internal representations (Pouget *et al*. [2000]; Panzeri *et al*. [2015]). For example, in fMRI research, activation-based methods are often used to infer that a brain region is involved in some mental process given its engagement during an experimental condition. But as a dependent measure, mean BOLD activity itself likely has no obvious functional significance. Similarly, the evoked responses that are the focus of traditional EEG and MEG analysis are not signals that the brain itself processes. In contrast, if the brain uses population codes, searching for information in patterns of activation means looking for the currency in which the brain makes its transactions.

As an illustration of the informational benefits of MVPA over univariate methods, consider the early findings of Haxby *et al*. ([2001]). Traditional univariate methods had previously been used to isolate the ‘fusiform face area’ (FFA) within the temporal cortex, which had been interpreted as a highly specialized face-processing ‘module’ in the ventral visual stream (Kanwisher *et al*. [1997]). Haxby *et al*. used MVPA to show that face information was discriminable in the ventral stream even when FFA was removed from the analysed ROI. Hence, their results demonstrated that decoding methods could reveal information present in brain activity that was otherwise undetectable by traditional methods. The results of Haxby *et al*. not only illustrated the greater sensitivity of decoding methods, but also made explicit the idea that decoding was potentially useful for revealing distributed representations in the brain.

In summary, univariate ‘activation-based’ analyses often obscure the information latent in spatial patterns of neural activity, while decoding affords a powerful tool for revealing this information. If the brain uses population codes, then spatial patterns in neural data that differentiate between conditions should be recoverable using information-based MVPA methods.

## 3. Why The Decoder’s Dictum Is False

Significant decoding indicates that information is latent in patterns of neural activity. However, researchers often draw a further inference: if there is decodable information, then there is strong evidence that the information is represented by the patterns of activity used as the basis for the decoding.

For example, Kriegeskorte and Bandettini ([2007], p. 658) claim that information-based analyses including MVPA ‘can help us look into [brain] regions and illuminate their representational content’. and go so far as to define decoding as ‘the reading out of representational content from measured activity’ (p. 659). Similarly, in comparing and contrasting different fMRI analysis techniques, Davis and Poldrack ([2013], p. 120) state that ‘[w]hereas univariate analysis focuses on differences in mean signal across regions of cortex, MVPA focuses on the informational content of activation patterns coded in different regions’. We have dubbed this further inference the Decoder’s Dictum. Although the Dictum is commonplace, exceptions can be found where decodability is observed but the interpretation of the results does not reflect this problematic inference. Instead, decodability is taken as evidence of functionally specialized processing rather than representational content (Davis and Poldrack [2013]).

The many fMRI decoding studies looking at top-down effects of visual and cognitive processing on primary visual cortex (V1) provide a good illustration. For example, Williams *et al*. ([2008]) presented simple object exemplars in the visual periphery, and found that object shape could be decoded from foveal V1. Jehee *et al*. ([2011]) similarly found that if two orientation grating stimuli were presented in the periphery, but only one was attended to, this resulted in greater classification accuracy for the orientation of the attended stimulus. Both of these results were interpreted as providing evidence of attention-driven feedback to primary visual cortex. In another study, Kok *et al*. ([2012]) found that when the orientation of a grating corresponded with an observer’s expectations, this resulted in lower BOLD activity but higher classification accuracy. Again, the focus was on showing that early visual processing can be modulated by expectations. Finally, Harrison and Tong ([2009]) found that stimulus information in a working memory task could be decoded from V1 over a prolonged period of time, suggesting a recruitment of the region for preserving stimulus information for later recall. The common goal of these studies is to reveal facts about functional processing or localization, not representational content.

In what follows, we defend the strong claim that the Decoder’s Dictum is false: successful decoding of information does not provide reasonable grounds for the inference that patterns of neural activity represent the conditions (or aspects of the conditions) about which they carry information. For some philosophers, this might sound like a trivial point: of course we cannot make inferences from information to representation, as there is more to representation than merely carrying information. Fair enough. Yet the problem is not (just) that informational content comes too cheaply in comparison to representational content (Fodor [1984]). For even if we accept that neural representations have content that is partially, or wholly, determined by information, there are several reasons for thinking that the Dictum fails to hold. In the rest of this section, we argue that a fundamental methodological issue with MVPA—specifically, the uncertainty regarding the information exploited by linear classifiers—shows why the Dictum is false.

### 3.1 We don’t know what information is decoded

The Dictum entails that if a classifier can discriminate between conditions, then it is picking up on the same information encoded by underlying neural representations. The problem is that we rarely know what information a classifier actually relies on. Indeed, this is most obvious in cases where we know a good deal about what a brain region represents.

To illustrate, consider again V1, where we have a reasonably good understanding of how orientation information is encoded (see, for example, Priebe and Ferster [2012]). Orientation-related information is also highly decodable using fMRI and MVPA (Haynes and Rees [2005]; Kamitani and Tong [2005]). And yet, we do not know what information classifiers are extracting from this region. Indeed, it is something of a mystery why fMRI decoding in the region even works at all. A typical voxel during a functional scan has a much coarser spatial resolution (> 2 x 2 x 2 mm) than the scale of the cortical columns that code for orientation in this region (∼2 mm in humans; ∼ 1 mm in monkeys). This means that one plausible explanation about how decoding works—that patterns of activity across orientation columns occur at a spatial scale roughly commensurate with the resolution of fMRI—cannot be correct.

There are a number of competing hypotheses about how orientation decoding in V1 is possible. Imperfect sampling of the underlying orientation columns might result in small biases at the voxel level, which decoding exploits, resulting in ‘hyperacuity’ or sub-voxel resolution (Haynes and Rees [2005]; Kamitani and Tong [2005]). Another possibility is that biases in the retinotopic map in V1 (in particular, radial biases) enable successful orientation decoding (Mannion *et al*. [2009]; Freeman *et al*. [2011]). Yet a third possibility is that activity patterns elicited by stimulus edges, not sampling or retinotopic biases, provide a potential source of decodable information in V1 (Carlson [2014]). Note here that the ‘biases’ appealed to in the explanations of orientation decoding are (in some important sense) artifacts in the way the data presented to the classifier is structured, rather than deep facts about the representational structure of the brain. So long as there is any information that distinguishes the conditions at hand, a linear decoder stands a good chance of finding it.

These issues are not restricted to decoding orientation in V1. For instance, it has been found that motion information decoding is more robust in V1 than V5/MT+ (Kamitani and Tong [2006]; Seymour *et al*. [2009]). This result is surprising when one considers that the majority of MT+ cells encode motion direction, while < 50 % of V1 neurons exhibit motion sensitivity and the region does not have cortical columns for motion direction as it does for orientation (Lu *et al*. [2010]). Wang *et al*. ([2014]) observe a direction-selective response bias that appears to explain this contrast between decoding and underlying functional organization—it is present in V1-V3 but not in MT+—suggesting that motion decoding in early visual cortex bares little relation to the actual encoding structure of these regions.

Thus, the fact that decoding can pick up on information unused by the brain, even in regions where there is a suitable representation that is used (for example, orientation representation in V1), means that even when prior theory and decoding are in agreement, decoding results cannot be reliably interpreted as picking up on the information that is neurally represented and used. All the worse, then, when we do not have converging evidence and prior theory. This epistemic uncertainty regarding the source of decodable information cuts to the core of the theoretical rationale for the Dictum. It is for this reason it is false, as we will illustrate by reconstructing the theoretical basis for the Dictum. Although appeals to the Dictum are commonplace in research using MVPA (a point we will return to), the theoretical basis for the Dictum is often underspecified. Here we reconstruct the rationale. Doing so demonstrates why epistemic uncertainty regarding the source of decodable information is fatal for the Dictum.

### 3.2 The theoretical basis for the dictum

The Decoder’s Dictum licenses inferences from decodability to facts about neural representation. The principle is evidential: if we can decode, we have reasonably strong evidence about what is represented in the measured patterns of neural activity. But why think the Dictum is true? Here we reconstruct what we take to be the underlying theoretical basis for the Dictum.

The support for the Dictum starts with two seemingly uncontroversial claims. The first is that if activity patterns occurring in different experimental conditions are discriminable, then information about the conditions is latent in these patterns. The second is that if activity patterns represent information about an experimental condition, then there must be some way to decode that content from the neural patterns. In other words, if internal representations are implemented in patterns of neural activity, and the brain is an encoder and decoder of its own neural signals, then the information must be decodable—that is, after all, what makes it a code. While substantive, these assumptions are not enough to get us to the Dictum. For all we have said, representations present in the brain might not have the right relationship to information extracted by MVPA when applied to the data recorded with standard neuroimaging techniques.

Two additional steps are required. The first secures the link between information and representation. This requires something like an informational approach to internal representations and their content. The presence of a statistical dependency or correlation is of interest because it suggests a causal dependency between the patterns and the experimental conditions (cf. Dretske [1983]). So charitably, the notion of information that researchers have in mind is that of natural information, where an event caries natural information about events that reliably cause it to occur (Scarantino and Piccinini [2010]). The view, which many in the field endorse, is very similar to Dretske’s ([1988]): a representation is a state that carries natural information, appropriately formatted to function as a state carrying this information.

For example, Cox ([2014], p. 189) notes that decoding research on the visual system:

> implicitly recognizes that the problem of vision is not one of information content, but of format. We know that the activity of retinal ganglion cells contains all of the information that the visual system can act upon, and that nonlinearity and noise in neuronal processing can only decrease (and never increase) the absolute amount of information present. However, the information present in the firing of retinal ganglion cells is not in a format that can be easily read-out by a downstream neuron in order to guide action.

In other words, vision repackages the information latent in the retinal input to make it functionally available for downstream perceptual and cognitive processing. A simple informational theory of representational content has as a corollary the idea that we can distinguish between implicit and explicit information (Kirsh [1990]), where being ‘implicit’ or ‘explicit’ is understood as being relative to some procedure for reading-out the information based on how a code is structured. Why should we think that successful decoding allows us to make an inference about what information is explicitly represented by a population code? This question brings us to the second additional assumption: the biological plausibility of MVPA methods in general, and linear classifiers in particular.

Many views of population coding assume that information can be read out by some sort of linear combination of components to the code. If so, then properties of the code can be made salient in the appropriate activation space. As Kriegeskorte and Kievet ([2013], p. 401) put it:

> We interpret neuronal activity as serving the function of representing content, and of transforming representations of content, with the ultimate objective to produce successful behaviors [⋯] The population of neurons within an area is thought to jointly represent the content in what is called a neuronal population code. It is the pattern of activity across neurons that represents the content [⋯] We can think of a brain region’s representation as a multidimensional space [⋯] It is the geometry of these points that defines the nature of the representation.

Now comes the crucial step. If population coding does indeed involve linear combination of elements, then MVPA is a plausible way to extract that information. For ultimately, a linear classifier is a biologically plausible yet abstract approximation of what the brain itself does when decoding its own signals (DiCarlo and Cox [2007]; King and Dehaene [2014]). In other words, because of the biological plausibility of linear classifiers, significant decodability is taken as evidence that the latent information in the data is also explicitly represented in the brain.

It is explicitly assumed in the field that linear decodability suffices to reveal an explicit representation. In fact, Kriegeskorte and Kievet ([2013], p. 402) go so far as to define explicit representation in such terms, claiming that ‘if the property can be read out by means of a linear combination of the activities of the neurons [⋯] the property is explicitly represented.’

Misaki *et al*. ([2010], p. 116) offer a similar characterization of when information is explicit:

> Linear decodable information can be thought of as “explicit” in the sense of being amenable to biologically plausible readout in a single step (i.e. by a single unit receiving the pattern as input) [⋯] Linearly decodable information is *directly* available information [⋯]

So the decoding of a linear classifier serves as a surrogate for the decoding of the brain. If the linear classifier can use information latent in neural activity, then this information must be used (or usable) by the brain: decoding provides evidence of an encoding.

In summary, one gets to the Decoder’s Dictum by endorsing several claims: (1) that MVPA reveals information latent in neural activity; (2) that an underlying neural population code implies decodability; (3) an informational view of neural representations and their contents; and (4) the hypothesis that biologically plausible linear classifiers are sufficiently similar in architecture to the decoding procedures employed by the brain. The latter is what lets us infer that decodable information is appropriately formatted for use by the brain, even when we do not necessarily know what that format is. So (5): if we can decode information from patterns of activity using MVPA, this provides good evidence in favor of the hypothesis that the patterns represent the information. Which is just a restatement of the Dictum.

### 3.3 Undermining the theoretical basis

We are now in a position to see precisely why the Dictum is false. For starters, note that a version of the Dictum appealing to nonlinear classifiers would be summarily rejected by researchers, as one cannot make an inference about what information is represented by patterns of neural activity using overpowered, biologically implausible nonlinear methods. For example, Kamitani and Tong ([2005], p. 684) were the first to caution against the use of nonlinear classifiers:

> [⋯] nonlinear methods may spuriously reflect the feature-tuning properties of the pattern analysis algorithm rather than the tuning properties of individual units within the brain. For these reasons, it is important to restrict the flexibility of pattern analysis methods when measuring ensemble feature selectivity.

Along the same lines, Naselaris *et al*. ([2011]) point out that nonlinearity should be avoided precisely because it is too powerful: it allows us to pull out information that is present in the brain, but that could not be exploited by the brain itself. Hence even though:

> [i] n theory a sufficiently powerful nonlinear classifier could decode almost any arbitrary feature from the information contained implicitly within an ROI⋯a nonlinear classifier can produce significant classification even if the decoded features are not explicitly represented within the ROI. (Naselaris *et al*. [2011], p. 404).

The concern is that information relied on by nonlinear classifiers might bear little relationship to what is actually represented by the brain. In other words, nonlinear classifiers are too informationally greedy, and so cannot serve as surrogates for the decoding procedures of the brain. Hence, a version of the Dictum appealing to nonlinear classifiers would clearly be false: nonlinear decoding does not provide evidence for what neural activity patterns represent. In contrast, the standard version of the Dictum seems to assume that linear classifiers are relatively conservative in terms of the information they can exploit (that is, they are biologically plausible), and so provide a safe (if defeasible) basis for making claims about representational content. The fact that a linear classifier can discriminate between activity patterns from different conditions is taken to provide good evidence that information about the conditions is both latent in the brain and functionally available.

Critically, our earlier discussion of the uncertainty surrounding the source of (linearly) decodable information shows the flaw in this reasoning. The fact that linear classifiers are biologically plausible does not preclude them from also being informationally greedy. Linear classifiers are surprisingly good at finding some linear combination of input features which discriminates between conditions in a multivariate data set. As we saw in our discussion of orientation decoding in V1, even when we do know the underlying functional architecture, how a classifier exploits information in neural data is deeply opaque. To further illustrate the greed of linear classifiers, consider that in psychology some have noted that linear decision-making models can be surprisingly good even when feature weightings are assigned more or less arbitrarily (Dawes [1979]). To emphasise a similar point, when using MVPA there is not even a guarantee that classifiers are detecting multivariate signals. In a simulation study, Davis *et al*. ([2014]) produced a univariate fMRI signal that could not be detected by activation-based analyses, but could nonetheless be decoded reliably.

Although a classifier (linear or nonlinear) may, through training, come to discriminate successfully between activity patterns associated with different experimental conditions, the information the classifier uses as the basis for this discrimination is not constrained to be the information the brain actually exploits to make the distinction (that is, they are informationally greedy). Importantly, it is evidence about the latter and not the former that is critical for zeroing in on the contents of neural representations. Hence, decodability does not entail that the features being combined, or their method of combination, bears any connection to how the brain is decoding its own signals. At best, MVPA-based decoding shows that information about experimental conditions is latent in neural patterns, but it cannot show that it is used, or even usable, by the brain. This is the deep reason why the Dictum is false.

## 4. Objections And Replies

We have argued that the Decoder’s Dictum is false. In this section we consider and respond to some objections to our criticism.

### 4.1 Does anyone really believe the Dictum?

When criticizing inferences in cognitive neuroscience, it is common for the philosopher to be informed that no working scientist really makes the sort of inference. Such an assertion is often meant to be a normative claim as much as a descriptive one (‘no good scientist argues thus’). Yet it is the descriptive claim which really matters—for philosophical critique matters only insofar as it identifies areas of actual methodological friction.

Do scientists really believe something like the Dictum? Our reconstruction of the theoretical basis of the Dictum already suggests that they do. At the same time, enumeration is also illuminating. Here are just a few (of many possible) illustrative examples where the Dictum is either overtly referenced or strongly implied:

1. Kamitani and Tong ([2005]) was one of the first studies showing that orientation information is decodable from voxels in early visual cortex, including V1. They state that their MVPA approach ‘may be extended to studying the neural basis of many types of mental content’ (p. 684).
2. Hung *et al*. ([2005]) was one of the first studies to pair MVPA with cellular recordings. They showed that object identity and category could be decoded from monkey IT as soon as ∼125 ms post-stimulus onset. They state that their approach ‘can be used to characterize the information represented in a cortical area [⋯]’ (p. 865).
3. In an early review of studies like Kamitani and Tong ([2005]) and Hung *et al*. ([2005]), Haynes and Rees ([2006], p. 524) conclude that ‘individual introspective mental events can be tracked from brain activity at individual locations when the underlying neural representations are well separated’, where separation is established by decodability with linear classifiers.
4. Woolgar *et al*. ([2011]) used decoding to investigate the multiple-demand or ‘MD’ regions of the brain, a frontoparietal network of regions that seem to be recruited across cognitive tasks. They used decoding to investigate these regions because ‘[i]n conventional fMRI the representational content of MD regions has been more difficult to determine, but the question can be examined through multi-voxel pattern analysis (MVPA)’ (p. 744).
5. An important technique with time-series decoding is that of discriminant cross-training, or ‘temporal generalization’: a classifier is trained on data from one time-bin, and tested on another. In a review of this method, King and Dehaene ([2014], p. 1) claim it ‘provides a novel way to understand how mental representations are manipulated and transformed’.
6. More complex MVPA methods, which characterize the structure of an activation space, or its ‘representational geometry’, have been promoted as ‘a useful intermediate level of description, capturing both the information represented in neuronal population code and the format in which it is represented’ (Kriegeskorte and Kievet [2013], p. 401).

Some brief observations are worth making about these examples. First, they include both individual studies (1, 2, 4) and reviews (3, 5, 6), spanning most of the period that decoding methods have been utilized in neuroimaging, and were written by key figures responsible for developing these methods. Second, the examples span fMRI (1, 4), EEG and MEG (4), and cellular recordings (2, 3). The Dictum thus appears to be a fundamental and widespread assumption in cognitive neuroscience, which has arguably played a key role in popularizing MVPA because of what it promises to deliver.^5^

### 4.2 Good decoding is not enough

Another tempting reply to our argument goes as follows. Classifier performance is graded, so it makes sense to talk about different brain regions having more or less decodable information. For example, although early visual cortex contains some information about object category, decodability is typically much worse than it is in inferior temporal cortex (IT), a region heavily implicated in the representation of object categories (Kiani *et al*. [2007]; Kriegeskorte *et al*. [2008]). So perhaps the Dictum is true if we restrict ourselves to the best or most decodable regions.

The problem with this reply is that it faces the same objection elaborated in detail above. What makes a given region the best or most decodable might have little or nothing to do with the information that is available to and used by the brain. This is why decoding results can be (and often are) at odds with the answers derived from other methods. As pointed out earlier, visual motion is more decodable from V1 than V5/+MT using fMRI (Kamitani and Tong [2006]; Seymour *et al*. [2009]), even though it is well-established that V5/+MT is a functionally specialized region for representing and processing motion information. Seymour *et al*. ([2009]) similarly report classification accuracy of 86 *%* in V1 and 65 *%* in V5/+MT, though they themselves refrain from drawing any strong conclusions due to the ‘potential differences underlying functional architecture in each region’ (Seymour *et al*. [2009], p. 178).

Their caution appears to embody the same concern that decoding results may reflect arbitrary differences to which the classifier is sensitive, without guaranteeing that these results track real differences in neural representation. Decoding—excellent or otherwise—is not a reliable guide to representation.

Another problem with this suggestion is that it entails that poor decodability (or even failure to decode) provides evidence that the information is not represented in a region. But this is false. Non-significant decoding does not entail the absence of information. One might have simply chosen the wrong classifier or stimuli, or the particular code used by the brain might not be read out easily by a linear classifier. Dubois *et al*. ([2015]) provide a nice illustration of this issue. They compared single-unit recordings with fMRI decoding in the face patch system of the macaque brain—an area known to possess face-sensitive neurons. In agreement with the single-unit data, face viewpoint was readily decodable from these regions. However, in the anterior face patches, face identity could not be decoded, even though single unit data shows that it is strongly represented in the region. These results indicate how poor decodability provides a thin basis upon which to mount negative claims about what a given region does not represent.

In sum, one cannot appeal to any level of classifier performance—good or bad—to preserve the Dictum.

### 4.3 Predicting behaviour is not enough

Though not always carried out, the ability to connect classifier performance to behaviour has been highlighted as one of the strengths of decoding methods (Naselaris *et al*. [2011]). To be sure, a deep problem with the Dictum is that decodability fails to show that information is formatted in a way that is used, or usable, by the brain (Cox and Savoy [2003]), while connecting decoding to behaviour helps make the case for functional utilization (Tong and Pratte [2012]). If behavioural performance can be predicted from the structure present in brain activation patterns, this would provide more compelling evidence that decodable information is used (or at the very least usable) by the brain, and hence neurally represented.

The simplest way to connect decoding and behaviour is to show that classifier and human performance are highly correlated. Minimally, if this obtains for some activation patterns more than others, this provides some (relatively weak) evidence that the patterns which correlate with behaviour represents information that is used in the guidance of behaviour.

Williams *et al*. ([2007]) provided one of the earliest indications that not all decodable information is ‘read-out’ in behaviour. They analysed the spatial pattern of the fMRI response in specific task-relevant brain regions while subjects performed a visual shape discrimination task. They hypothesized that if decodable shape category information is behaviourally relevant, then decodability should be higher on correct trials than on incorrect trials. Critically, they showed that although both retinotopic cortex and lateral occipital cortex (LOC) in humans contains decodable category information, only the LOC shows a difference in pattern strength for correct as compared to incorrect trials. Specifically, category information was decodable on correct but not incorrect trials in the LOC. This was not true for retinotopic cortex. This pattern of results suggests that only the information in LOC might drive behaviour.

It is also possible to quantify the relationship between decodability and behaviour more precisely. For example, in an early EEG decoding study, Philiastides and Sajda ([2006]) were able to show there was no significant difference between human psychometric and classifier ‘neurometric’ functions, suggesting that the classifier performance was highly predictive of observer performance when trained on time-series data of certain latencies.

While connection to behaviour supplies valuable evidence, we still think that it is not enough to warrant inferences to representational content. As we noted earlier, there are cases where decodability appears to show something about functional processing rather than the content of neural representations. Again, V1 provides a useful test case. Since we know that V1 primarily encodes information about low-level visual features (such as luminance or orientation) and does not encode higher-level visual features (such as shape or object category) any decoding of higher-level visual features is unlikely to reflect genuine representational content. This is true even if decoded information can be linked with behavioural performance. For example, Haynes and Rees ([2005]) found that V1 activity was predictive of whether or not subjects were perceiving a visual illusion, and Kok *et al*. ([2012]) found that top-down effects of expectation on V1 were predictive of behavioural performance. In these cases, the connection is that early processing modulates later processing that determines behaviour.

Note that the problem is not one of spurious correlation. In an important sense, it is quite the opposite problem. There is plenty of information, even in V1, which a clever decoding algorithm can often pick up on. More generally, a brain region might carry information which is reliably correlated with the information that is actually used, but which is not itself used in behaviour. This is because the information in a region might need to be transformed into a more appropriate format before it is read out. As DiCarlo and Cox ([2007], p. 335) put it, ‘[⋯] the problem is typically not a lack of information or noisy information, but that the information is badly formatted[⋯]’. But even ‘badly formatted’ information might correlate with behaviour. In summary, merely predicting behaviour using decodable information is not enough to revive the Dictum.

## 5. Moving Beyond The Dictum

We have argued that the Decoder’s Dictum is false. However, we are not pessimists about decoding. Rather, we think the right conclusion to draw is that decoding must be augmented in order to provide good evidence about neural representation. If linear classifiers are greedy, then they cannot function as a surrogate for the sort of linear read-out carried out by the brain. Instead, we need some additional assurance that a particular decoding result relies on information stemming from neural representations. This need not be knock-down evidence, but decodability alone is not enough to do the job (as the Dictum suggests).

In the previous section, we considered one form of augmentation—linking decoding results to behavioural outcomes—and argued that it was insufficient. The problem was that linkages to behaviour do not show that the information is actually formatted in a useable way. Framing it this way, however, already suggests a solution. The Dictum relies on the idea that the biological plausibility of linear classifiers allows them to function as a kind of surrogate—the classifier-as-decoder takes the place of the brain-as-decoder in showing that information that is latent in neural activity is used, or usable (cf. de Wit *et al*. [2016]). We have shown that it cannot play this role. But if the information latent in patterns of neural activity can be used to predict observer behaviour based on a psychological model, then we would have a more sound evidential basis for drawing conclusions about neural representation. For unlike classifier performance, observer behaviour is clearly dependent on how the brain decodes its own signals. In other words, this approach depends on offering a psychologically plausible model of how observers (through down-stream processing) exploit the information found in patterns of neural activity (cf. Ritchie and Carlson [2016]). And as it happens, such an approach is already on offer.

There is a long tradition in psychology of modeling behavioural performance using psychological spaces (Attneave [1950]; Shepard [1964]). Here by ‘psychological’ space we mean a space in which dimensions reflect different features or combinations of features of stimuli, as reconstructed from comparative similarity judgments of observers of stimuli/conditions. Models within this tradition characterize representations for individual stimuli or experimental conditions as points in a space, and observer behaviour (such as choice or reaction time) is modeled based on the relationship between different representations in these spaces. Thus, familiar categorization models from cognitive psychology such as prototype models, exemplar models, and decision boundary models all predict observer behaviour based on different distance metrics applied to a reconstructed psychological space (Ashby and Maddox [1993]). A virtue of some MVPA methods like Representational Similarity Analysis (RSA) is that they help to focus attention on structure in activation spaces (Haxby *et al*. [2014]; Kriegeskorte and Kievet [2013]). In RSA the pair-wise (dis)similarity for patterns of activity for different conditions is computed, which can be used to reconstruct an activation space from multivariate neural data. A hypothesis that many have considered is that if an activation space implements a psychological space, then one can apply psychological models or hypotheses to the activation space directly in order to predict behaviour (Edelman *et al*. [1998]; de Beeck *et al*. [2001], [2008]; Davis and Poldrack [2014]). Note that this approach is importantly different from the Dictum, as it does not rely on using linear classifiers as a surrogate. Furthermore, the approach achieves both biological and psychological plausibility through a linkage between the structure of the decoded activation space and the structure of behaviour (Ritchie and Carlson [2016]). And since it makes use of MVPA in conjunction with established techniques for modeling behaviour, it also takes advantage of some of the strengths of MVPA we have already mentioned. Here we offer two examples of research that adopt this sort of approach.

First, a popular and theoretically simple approach involves directly comparing the similarity structure of activation spaces with psychological spaces reconstructed from subjects’ similarity judgments of stimuli (e.g. Mur *et al*. [2013]; Bracci and de Beeck [2016]; Wardle *et al*. [2016]). One illustration of this approach is provided by the results of Sha *et al*. ([2015]), who collected similarity ratings for a large number of exemplar images for several animate or inanimate object categories. The similarity space constructed from these judgments was then directly related to the similarity structure of activation spaces from throughout the brain measured using fMRI. They found that activation spaces that correlated with the behavioural similarity space were best accounted for by a single dimension, which seemed to reflect an animacy continuum rather than a categorical difference between the neural patterns for animate and inanimate objects (Kiani *et al*. [2007]; Kriegeskorte *et al*. [2008]).

Second, some work has focused on the psychological plausibility of activation spaces by using them to predict the latency of behaviour. For example, in two studies using fMRI and MEG decoding, Carlson and Ritchie (Carlson *et al*. [2014]; Ritchie, Tovar, and Carlson [2015]) showed that distance from a decision boundary for a classifier through activation space was predictive of reaction time (RT). In their experiments they were explicitly motivated by the idea that linear classifiers are structurally identical to the model of an observer under signal detection theory (Green and Swets [1966]). A natural extension of signal detection theory is that distance from an evidential boundary negatively correlates with RT (Ashby and Maddox [1994]). As predicted, they found that RT negatively correlated with distance from the decision boundaries, suggesting a level of psychological plausibility to even simple linear classifiers.

Crucially, in these sorts of studies it is implausible to suppose that the information is present but not correctly formatted, because the decoded format of the information in activation space is precisely what is being used to predict behaviour in a psychologically plausible manner. We do not mean to suggest that the results we have summarized suffice for drawing conclusions about neural representation, but we do believe that they help point the way forward.

## 6. Conclusion

The Decoder’s Dictum is false. Significant decoding, even when supplemented by other results, does not warrant an inference that the decoded information is represented. However, we do believe that if behaviour can be connected to the structure of activation space in a psychologically plausible manner, then this may warrant the sort of inference researchers have had in mind. And we should stress that we do not think the above shows that decoding is immaterial. Indeed, as we have suggested, MVPA is crucial for connecting activation spaces to behaviour. Rather, as we have argued, appealing to the Dictum obscures not only the true import of decoding as a tool in cognitive neuroscience, but also what sort of evidence is required for making claims about neural representation.

## Acknowledgements

Thanks to two anonymous reviewers for helpful comments on a previous draft, and to audiences at Macquarie University and the 2014 Australasian Society for Cognitive Science for feedback on earlier versions of this work. Funding for this research was provided by the Australian Research Council (FT140100422 to Colin Klein).

## Footnotes

1 A recent example: in arguing against the encapsulation of the visual system, Ogilvie and Carruthers ([2016]) rely almost exclusively on decoding results about early vision since they believe it provides more convincing evidence than behavioural research.

2 One may reasonably wonder whether this characterization captures scientific usage. Although foundational concepts like ‘representation’ are rarely explicitly defined by neuroscientists, there are exceptions. For example, Marr ([1982], pp. 20-1) defines a representation as ‘a formal system for making explicit certain entities or types of information’, and Eliasmith and Anderson ([2003], p. 5) state that: ‘[r]epresentations, broadly speaking, serve to relate the internal state of the animal to its environment; they are often said to “stand-in for” some external state of affairs.’ Along similar lines, deCharms and Zador ([2000], p. 614) define a representation as a ‘message that uses [⋯] rules to carry information’ and define content as the ‘information that a representation carries’. Our discussion of the theoretical basis for the Dictum (section 3.2) also illustrates that something close to the above notion is widely assumed by researchers in the field.

3 Some terminological points. First, ‘MVPA’ originally meant ‘multi-voxel pattern analysis’, rather than *‘multivariate* pattern analysis’. The latter is preferable because it highlights the fact that the methods are not specific to fMRI (Haxby [2012]). Second, ‘MVPA’ and ‘decoding’ are sometimes used interchangeably (as we do), but strictly speaking decoding methods are a subset of MVPA methods (Naselaris *et al*. [2011]). And third, ‘decoding’ is often used in two distinct senses: a machine learning sense, in which it is basically a synonym for ‘classify’; and a neural sense, referencing the encoding and decoding of signals by the brain. We make use of both senses here.

4 It is well-known that the signals measured with neuroimaging techniques such as fMRI and MEG/EEG depend on neural activity, but often in complicated and indirect ways (e.g., Logothetis [2008]; Nir *et al*. [2008]; Singh [2012]). For example, fMRI measures blood oxygenation level-dependent (BOLD) signals reflecting changes in cerebral blood flow (CBF), cerebral blood volume (CBV), and cerebral metabolic rate of oxygen consumption (CMRO2) following neural activity. Although it remains controversial precisely which types of neural responses induce these haemodynamic changes (e.g., Logothetis *et al*. [2001]; Sirotin and Das [2009]; Lee et al. [2010]), applications of MVPA typically assume that neuroimaging techniques coarsely measure the spatial structure and temporal dynamics of local neuronal populations. It is therefore common to use the term ‘activity patterns’ to describe the multivariate data collected with these techniques, even though, strictly speaking, MVPA is not being used to analyse neural activity patterns directly. We also adopt this convention.

5 Of course, not all researchers using MVPA subscribe to the Dictum. As we have acknowledged, some embrace decoding because of its benefits over more conventional analyses, without drawing unjustified inferences about representational content.

